# Metaecosystem dynamics drive community composition in experimental multilayered spatial networks

**DOI:** 10.1101/675256

**Authors:** Eric Harvey, Isabelle Gounand, Emanuel A. Fronhofer, Florian Altermatt

## Abstract

Cross-ecosystem subsidies are studied with a focus on resource exchange at local ecosystem boundaries. This perspective ignores regional dynamics that can emerge via constraints imposed by the landscape, potentially leading to spatially-dependent effects of subsidies and spatial feedbacks. Using miniaturized landscape analogues of river dendritic and terrestrial lattice spatial networks, we manipulated and studied resource exchange between the two whole networks. We found that community composition in dendritic networks depended on the resource pulse from the lattice network, with the strength of this effect declining in larger downstream patches. In turn, this spatially-dependent effect imposed constraints on the lattice network with populations in that network reaching higher densities when connected to more central patches in the dendritic network. Consequently, localized cross-ecosystem fluxes, and their respective effects on recipient ecosystems, must be studied in a perspective taking into account the explicit spatial configuration of the landscape.

**Statement of authorship:** EH, IG, EAF and FA designed the research; EH conducted the lab experiment with support from IG, EAF and FA, processed the experimental data with methodological developments from IG, and carried out the analysis of experimental data; all authors participated in results interpretation; EH wrote the first draft of the manuscript; All authors significantly contributed to further manuscript revisions.

## Introduction

The significance of cross-ecosystem subsidies in supporting recipient ecosystems is well recognized (Polis *et al.* 1997; Richardson & Sato 2015; Soininen *et al.* 2015). For instance, Fisher & Likens (1973) estimated that small streams obtain 75% of their total energy budget from terrestrial sources. Those cross-ecosystem subsidies can support complex communities (Polis & Hurd 1995), and sometimes lead to indirect bottom-up effects across ecosystems via spatial trophic dynamics (Knight *et al.* 2005; Bultman *et al.* 2014; Koel *et al.* 2019). Cross-ecosystem subsidy studies generally focus on resource exchange at a specific ecosystem boundary (Gounand *et al.* 2018). This localized perspective is ignoring regional scale dynamics that can emerge via dispersal and spatial feedbacks: the effect of landscape configuration on population-level (Altermatt & Fronhofer 2018) and community-level (Tscharntke *et al.* 2012; Tonkin *et al.* 2018b) processes are well-documented. Surprisingly, however, cross-ecosystem dynamics scaling from localized resource flows at ecosystem boundary to landscape-wide effects remain unstudied.

In a cross-ecosystem context, the landscape can be represented as two or more spatial networks embedded within one another and interacting through the exchange of cross-ecosystem resources (Mucha *et al.* 2010) (Figure 1). Those spatial networks have different structures and properties. For instance, in natural landscapes, dendritic river networks are connected with terrestrial spatial networks by the exchange of resources (organic matter, inorganic nutrients, Figure 1). Each network also undergoes its own internal spatial dynamics characterized by the movement of organisms through dispersal or foraging behaviors that are constrained by the shape of the network itself. This internal dynamic leads to intrinsic spatial variations in biodiversity within each network (Harvey & MacDougall 2014; Tonkin *et al.* 2018a).

**Figure 1.**
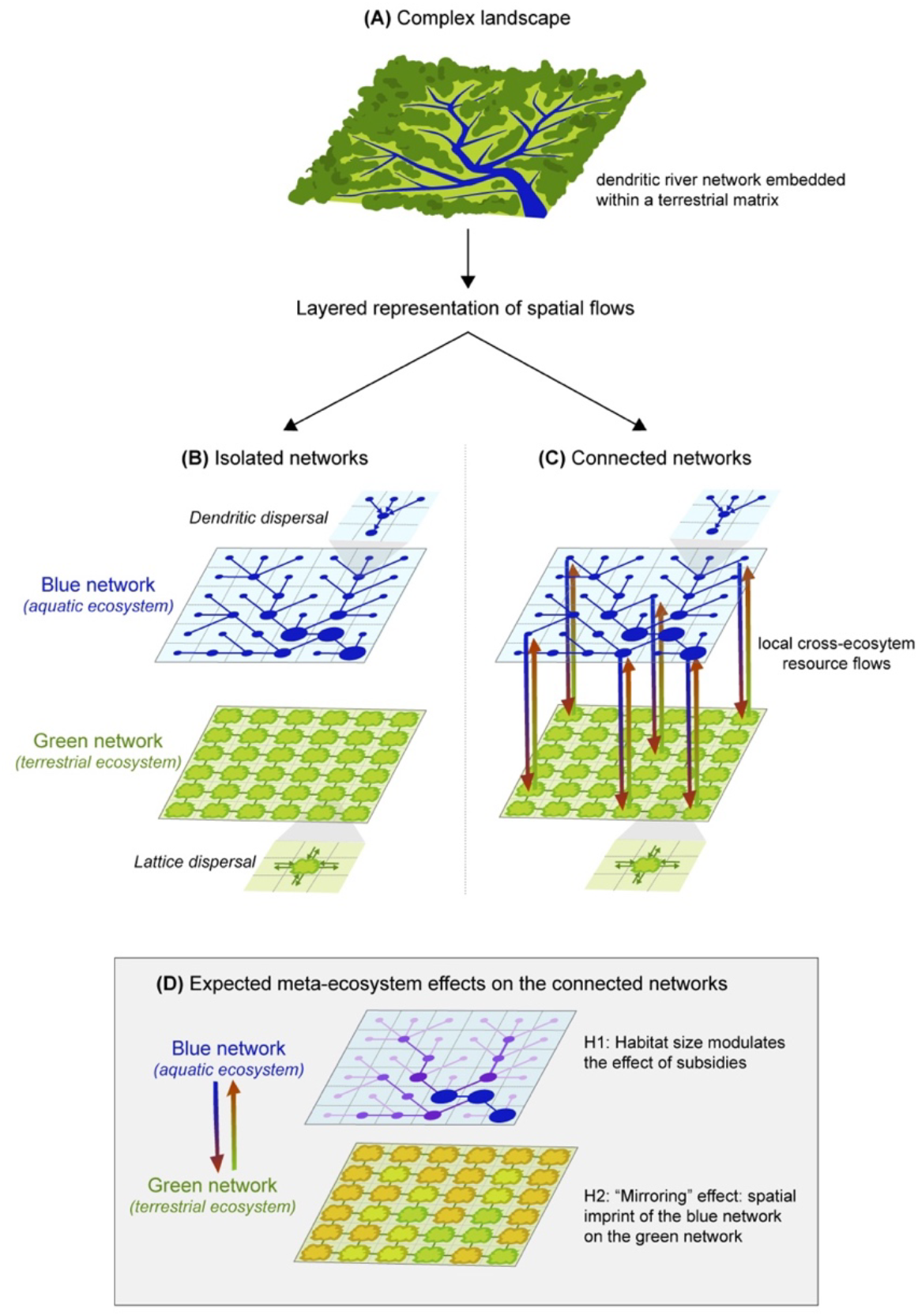
Conceptual representation of the experimental design. (**A**) Natural landscapes are composed of dendritic river networks embedded within terrestrial matrices. This complexity can be decomposed using a layered representation with two distinct spatial networks: an aquatic dendritic network and a terrestrial lattice network. (**B**) For each spatial network dispersal occurs along the edges, (**C**) while cross-ecosystem resource exchange is local and links each node in the respective position in the two networks (only few arrows drawn, for clarity). (**D**) Hypotheses on the interactive effects of cross-ecosystem resource exchange and spatial structure on green and blue networks

For instance, river networks are known to constrain alpha and beta diversity, generally leading to higher alpha and lower beta diversity in downstream than in upstream patches (Finn *et al.* 2011; Carrara *et al.* 2012; Tonkin *et al.* 2018a). Models and experiments have shown that those constraints on biodiversity can emerge from the topology of the dendritic network itself, which intrinsically drives the distribution of habitat capacity (i.e., volume or size) and dispersal limitation (Carrara *et al.* 2012). Assuming equal transfer of cross-ecosystem resource from the terrestrial network to the river network across the landscape, smaller and more isolated upstream patches should respond more to resource pulses than larger and more connected downstream patches, where the local effect of resource exchange is more likely to be diluted by larger volume. Smaller patches are also expected to be more resource-limited than larger ones, leading to a stronger dependency or response to external resource inputs (Polis & Hurd 1995). At the landscape scale, this would lead to spatial variation in the effects of cross-ecosystem resource exchange depending on the position in the river network.

Reciprocity in resource exchange between two spatial networks could also lead to spatially-dependent feedbacks where, for instance, effects from the terrestrial to the river network are especially important in upstream reaches (for reasons described above), in turn, leading to effects on the terrestrial network being especially important for terrestrial patches connected to upstream sites in the river network (Figure 1). Thus, via bi-directional cross-ecosystem resource exchange, one of the spatial networks can potentially impose their own structural constraints on the connected network leading to a “mirroring effect” influencing the entire spatial network but only visible at landscape extent.

In this study, we tested **i**) the influence of the position in the landscape on the effects of cross-ecosystem resource exchange and, subsequently, **ii**) if and how spatial feedbacks emerge at the landscape scale, leading to a “mirroring effect” on community composition. To meet those two objectives, we used microcosms in a controlled laboratory experiment, building miniaturized spatial network analogues, reflecting the general spatial properties of natural landscapes. Specifically, we connected a dendritic spatial network (representative of riverine networks – “blue network”) to a 4-nearest neighbor lattice network (representative of a terrestrial matrix embedding the river network – “green network”) by the exchange of resources (Figure 1). Each network was composed of local ecosystems (i.e., each microcosm) connected by the dispersal of living organisms along the specific structure of the respective network (Figure 1). At the local scale, each of these local ecosystems was also linked to a local ecosystem in the other network (i.e., across the blue and green network) by the exchange of resource only (i.e., inorganic nutrients and dead biomass). Thus, living organisms were moving only within each network, while dead biomass also moved between the two networks. The blue network contained seven interacting bactivorous protist species while the green network contained bacterial communities. This disparity in biotic complexity between the two networks made the bi-directional effect of resource exchange more tractable to test our working hypotheses. We focus on the influence of resource pulses from the green network on protist community dynamic in the blue network and, in turn, on how the altered dynamics along the blue network might feedback and affect the spatial distribution of bacterial densities in the green network compared to isolated controls (“mirroring effect”, Figure 1). Our large-scale (504 microcosms) experiment replicated across entire landscape-analogues allowed us to test how the landscape *per se* constrains spatial variation in the effect of cross-ecosystem resource exchange and how it scales up to generate regional-level patterns.

## Methods

Each network (‘blue’ and ‘green’) was represented by 36 microcosms connected by dispersal along its edges (Figure 1). We had four replicates of each network connected by resource exchange (288 microcosms). Then, as controls, we had four isolated (not connected to a green network) replicates of the blue network (144 microcosms) and two isolated (not connected to a blue network) replicates of the green network (72 microcosms –spatially homogenous dispersal). In total, we had 504 microcosms. The experiment lasted 29 days with 5 sampling events.

For the green network, we used a square lattice network (Figure 1). This choice was justified by the many previous theoretical and empirical metacommunity studies using simplified lattice networks to approximate connectivity and dispersal based on Euclidean distances in many terrestrial ‘2D’ systems (Hanski & Gaggiotti 2004; Leibold *et al.* 2004; Holyoak *et al.* 2005).

For the blue network, our four replicates corresponded to four different realizations of dendritic networks generated from four different space-filling optimal channel networks (Rigon *et al.* 1993). Optimal channel networks are known to reproduce the scaling properties observed in river systems (Rinaldo *et al.* 2006; Carrara *et al.* 2014). They are built under the assumption that drainage network configurations should minimize total energy dissipation, and the empirical observation that river network properties constitute scale-invariant fractals (Rinaldo *et al.* 2006). To reduce the four networks generated this way (corresponding to our four replicates for the blue network) to a logistically possible level for a laboratory experiment, a coarse-graining procedure was used to generate 6×6 patch networks of four different volumes (7.5, 13, 22.5 and 45 mL), preserving the characteristics of the original three-dimensional basins (Rodriguez-Iturbe & Rinaldo 1997; Carrara *et al.* 2014).

Biotic communities in the blue network were composed of six bacterivorous protist and one rotifer species (henceforth called “protists”): *Tetrahymena* sp. (Tet), *Paramecium caudatum* (Pca), *Colpidium striatum* (Col), *Spirostomum* sp. (Spi), and *Chilomonas* sp. (Chi), *Blepharisma* sp. (Ble) and the rotifer *Cephalodella* sp. (Rot). The latter two species can also to a lesser degree predate on smaller protists. The protists were feeding on a common pool of bacteria (*Serratia fonticola*, *Bacillus subtilis* and *Brevibacillus brevis*). Prior to the beginning of the experiment, each protist species was grown in monoculture in a solution of pre-autoclaved standard protist pellet medium (Carolina Biological Supply, Burlington NC, USA, 0.46 g protist pellets 1 L^−1^ tap water) and 10% bacteria inoculum, until they reached carrying capacity (for methodological details and protocols see Altermatt *et al.* 2015).

Each ecosystem in the green network was set at 10 mL. Biotic communities in the green network were composed of three bacteria species (*Serratia fonticola*, *Bacillus subtilis* and *Brevibacillus brevis*). It is noteworthy that we initially inoculated the green network also with an autotrophic protist species (*Euglena gracilis*). However, the species did not establish well in the network and all individuals died before or very soon after the start of the experiment. Because the species was inoculated at equal density in each ecosystem of the green network, we can safely assume that the death of all individuals did not generate significant within network variations in detritus (and considering the additional homogenizing effect of dispersal). For this reason, we assumed a zero-sum effect, and will not consider this species further.

Each ecosystem consisted of a 50 (blue networks) or 15 (green networks) mL polypropylene Falcon tube (VWR, Dietikon, Switzerland). At day 0, we pipetted an equal mixture of each of the seven protist species at carrying capacity into each ecosystem of the blue network to reach the corresponding volume (7.5 mL, 13 mL, 22.5 mL, 45 mL). Communities were allowed to grow 24 hours before the first dispersal event. Within-network dispersal and cross-ecosystem resource exchange occurred twice a week, while sampling of the communities for species count was done once a week (two dispersal/resource pulse events between each sampling with at least 48 hours between the last dispersal/resource pulse event and sampling). Sampling events and counting were done at day 0, 7, 15, 21, 29 of the experiment, while dispersal and cross-ecosystem resource pulse events occurred at day 1, 4, 8, 11, 16, 19, 22, 25 of the experiment. On the dispersal/resource pulse days, dispersal was always done first, so that the pulsed resource added to each patch would stay in that patch until the next dispersal event.

Dispersal was done by pipetting a fixed volume from one ecosystem to each of the connected ecosystems along the edges of the spatial network, using mirror networks (following methods developed in Carrara *et al.* 2012). We assumed higher dispersal in the blue (1 mL) compared to the green (0.5 mL) network to mimic the action of physical flows in riverine dendritic networks. Dispersal was bi-directional along each edge for both networks (e.g., 1 or 0.5 mL from ecosystem a to b and 1 or 0.5 mL from ecosystem b to a), which ensured the maintenance of the same volume in each ecosystem throughout the 29 days of the experiment. We implemented bi-directional dispersal to avoid the logistical challenge of maintaining equal ecosystem volumes. In a previous study, we conducted a sensitivity analysis including an extensive number of simulations to confirm that this experimental assumption on dispersal did not affect the network effects on species richness (see Appendix A and Figs. S2 & S3 in Harvey *et al.* 2018).

Cross-ecosystem resource pulse was done by exchanging dead biomass from one network to the other. First, a set volume was removed from each ecosystem in the blue (1 mL) and green (1.25 mL, see paragraph below for explanation on the volume difference) networks. Those volumes were then microwaved until boiling to turn all living cells into detritus (following methods developed in Harvey *et al.* 2016, 2017). After a cooling period, the microwaved samples were poured into the specific recipient ecosystem in the recipient network (see Figure 1). To control for the mortality effect, we performed the same steps of sampling and microwaving in the isolated control networks, with the difference that the microwaved volume was poured back to the ecosystem of origin.

At each measurement day, sampling was done by pipetting a total of 0.5 mL from each ecosystem of each network that was then used to measure bacteria (0.1 mL) and protist densities (0.4 mL). Removing 0.5 mL from microcosms in the blue network will have different impacts depending on ecosystem volume. For this reason, we compensated this volume lost on a weekly basis by exchanging 0.25 mL more volume from the green to the blue network (resource exchange is done 2 times/week, thus totally replacing the 0.5 mL). Protist abundance was measured by using a standardized video recording and analysis procedure (Pennekamp & Schtickzelle 2013; Pennekamp *et al.* 2015). In brief, a constant volume (34.4 μL) of each 0.4 mL sample was measured under a dissecting microscope connected to a camera for the recording of videos (5 s per video). Then, using the R-package bemovi (Pennekamp *et al.* 2015), we used an image processing software (ImageJ, National Institute of Health, USA) to extract the number of moving organisms per video frame along with a suite of different traits for each occurrence (e.g., speed, shape, size) that could then be used to filter out background movement noise (e.g., particles from the medium) and to identify species in a mixture (details were published in Appendix C of Harvey *et al.* 2018). Finally, for bacteria we measured densities using standard flow cytometry on fresh SYBR green fixated cells using a BD Accuri™ C6 cell counter (1/1000 dilution). For logistical reasons and because of time constraints, bacteria counts were only done for two of the four replicates in blue and green networks.

## Statistical analysis

The main objective of this experiment was to identify landscape-scale feedbacks between the two spatial networks connected by the pulse exchange of resources. Our focus was on the interaction term between position in the blue network and the resource pulse treatment. Our second main working hypothesis was the “mirroring” effect where we expected to find an imprint of the blue network within the green network (Figure 1).

### Effects from green to blue network

Our main response variable in the blue network was changes in protist community composition (i.e., abundance and occurrence) because it encompasses effects on both diversity and the more subtle influences on the structure and functioning of the community. To test for those changes in community composition, we used two complementary approaches: Redundancy Analysis (RDA) and log response ratio of the means (LRR). The RDA analysis was of the form C ~ E where C represented the Hellinger-transformed protist abundance community matrix and E the predictor matrix including the effects of resource pulse from the green network (main treatment), ecosystem volume in the blue network, closeness centrality (a measure of the number of steps required to access every other ecosystem from a given ecosystem in the network - *sensus* Freeman 1978) in the blue network, time (continuous experimental time), two-way interaction between resource pulse and ecosystem volume, and two-way interaction between resource pulse and time. We then ran a type 3 permutation ANOVA (999 permutations) to determine F-statistic and significance level for each term from the RDA at p < 0.05. Permutations in the ANOVA were stratified by each network replicate (the 4 network topologies) nested within a sampling day (discrete experimental time).

The RDA provided a general statistical test for the effect of the predictors of interest on protist community composition in the blue network, but no actual effect sizes. As a second complementary step, we explored the effect of resource pulse from the green network using log response ratio of the means (LRR). This approach served to confirm results from the RDA and more importantly allowed us to evaluate each protist species response and its effect size. The log response ratio of the mean was here defined as:

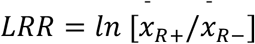

where ln is the natural log, 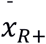 and 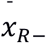 represent mean values (protist density) in the presence and absence of resource pulse, respectively. Following Hedges *et al.* (1999), the standard error of each LRR was calculated as:

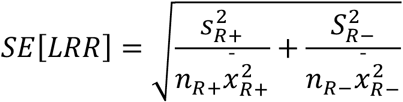

where *s* is the standard deviation and *n* is the sample size. Based on this measure of standard error we calculated confidence intervals (95%) for each LRR values. LRRs have straightforward interpretations: if the 95% confidence interval is not overlapping with 0, there is 95% probability that the population mean of effect size is indeed higher or lower than zero. Negative (or positive) LRR values means that the treatment, here resource pulse, had a negative (or positive) effect on population density.

As a complementary approach we also explored the effect of cross-ecosystem resource exchange on the community mean and standard deviation of protist traits related to body size. We used two traits measured during the video recording procedure and used for species identification: individual body length and width. Effects on community mean trait values, especially ones related to body size, can provide valuable insights on the mechanisms driving the observed changes in community composition and indicate potential, but unmeasured, impacts on ecosystem functions.

Finally, we also tested, using mixed-effect models, for effects of resource pulse from the green network on aggregate protist population and community metrics (total protist densities, species richness, evenness and bioarea) in the blue network. Evenness was measured as the Pielou’s evenness index (Pielou 1975). Protist bioarea per mL, used as a proxy of biomass, was calculated using the summed area of all individuals per video frame. The mixed effect models included the interactive effects of resource pulse, ecosystem volume and experimental time (continuous), and the effect of closeness centrality as a co-variate. To control for temporal pseudo-replication and measure the nonlinear variance associated with time (continuous time in the fixed model captures the linear trend), we added each network replicate nested within experimental day (discrete effect of time) as a nested random factor. The model was fitted by maximizing the restricted log-likelihood (‘REML’, see Pinheiro *et al.* 2018). For each model we decomposed the variation to evaluate the proportion of variance explained by the fixed terms and random factors.

### Effects from blue to green network

The green network was simpler in structure by design. Our main objective with the green network was to test whether we can detect spatial signals of the blue dendritic network on the green square lattice. Here we used a combination of a mixed-effect model and log response ratios of the means to test for the effect of resource pulse from the blue network, but also of the blue network structure and properties on bacteria density in the green network. Our main interest was to test whether bacteria density in a connected green ecosystem fluctuated depending on where in the blue network that ecosystem was connected (e.g., upstream vs. downstream). The global effect of resource pulse from the blue network was measured by LRR. Then as a second step, we used a mixed-effect model including the effects of ecosystem volume (in the blue network), closeness centrality (in the blue network), protist density, richness, and bioarea (in the blue network) on bacteria density in the connected green networks. To control for temporal pseudo-replication, we added experimental day (discrete effect of time) as a random factor. Again, the model was fitted by maximizing the restricted log-likelihood.

All analyses were conducted with R 3.5.1 (R Core Team 2018), using the ‘bemovi’ package (ver. 1.0) for video analyses (Pennekamp *et al.* 2015), the ‘vegan’ package (ver. 2.5-3) for multivariate analysis (Oksanen *et al.* 2018), the ‘nlme’ package (ver. 3.1-137) for the mixed-effect models (Pinheiro *et al.* 2018), the ‘igraph’ package (ver. 1.2.2) to extract network metrics (Csardi & Nepusz 2006), and the ‘ape’ (ver. 5.3) package to decompose the variation of each mixed effect models (Paradis & Schliep 2018).

## Results

Testing for landscape-scale effects of cross-ecosystem resource exchange between two distinct spatial networks, we found that protist community composition from the blue networks connected to a green network by resource pulse differed from communities in isolated blue networks (Table 1, Figure 2). Resource pulse from the green network also led to changes in trait values (Figure 2) that were predictable based on species body size and species-specific response to resource pulse (Figure 3). Changes at the population level scaled up to affect aggregate community metrics related to total protist density, richness, and evenness (see Table S1 in Supporting Information). The effect of resource pulse in the blue network also varied depending on the position in the network with strongest effects found in smaller upstream ecosystems (Table 1 and Figure 4). In turn, we found that bacteria density in the green network’s ecosystems was significantly higher when connected to a blue network (Figure 2), and this effect varied depending on how central was the ecosystem in the blue network that it was connected to (Table S2, Figure 5).

**Table 1.** Permutation ANOVA (999 permutations) on the Redundancy Analysis model used to test effects on protist community composition (abundance and occurrence) in the blue network.

|  | **Df** | **F** |
| --- | --- | --- |
| Resource pulse (R) | 1 | 49.97*** |
| Ecosystem volume (V) | 1 | 59.20*** |
| Centrality | 1 | 12.21*** |
| Time (T) | 1 | 237.65*** |
| R x V | 5 | 2.38*** |
| R x T | 5 | 15.65*** |
| Residuals | 1137 |  |

\*\*\* 0.001; \*\* 0.01; \* 0.05

**Figure 2.**
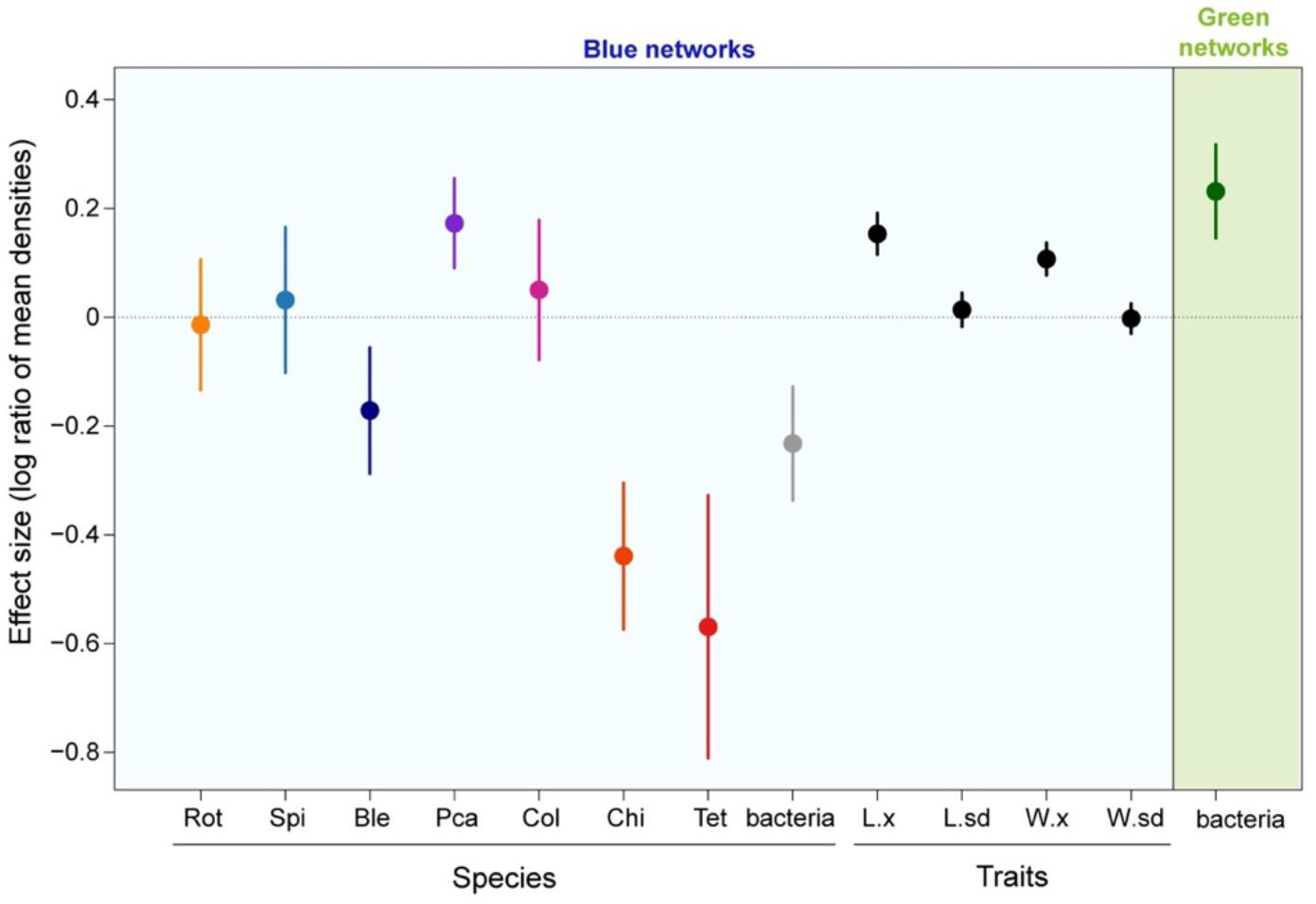
Effect size of the cross-ecosystem resource pulse respectively on protist taxa and bacteria density, and on community mean and standard variation of trait values in the blue network, and finally on bacteria density in the green network. Each point represents the log ratio of the mean effect of the treatment (as described in the Methods section). Error bars represent the 95% confidence interval calculated based on the LRR standard error formula described in the Methods section. An LRR higher than zero represents a positive effect of resource pulse on density and a negative LRR, the opposite. For body length (L) and width (W), x represents the mean and sd the standard variation of the mean trait value. The different protist taxa are color coded and named by their abbreviations described in the Methods section.

**Figure 3.**
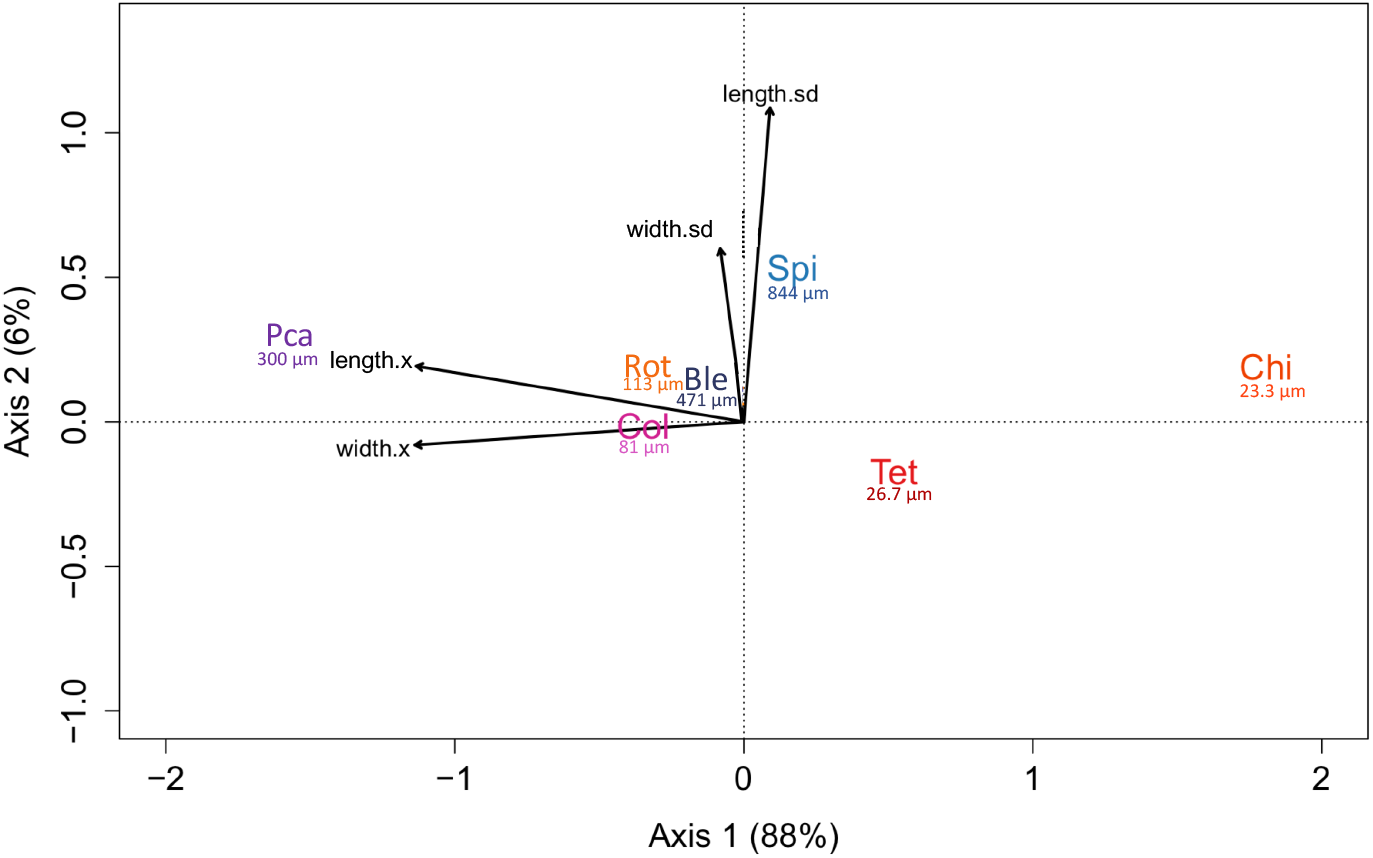
Redundancy analysis showing the association between community mean trait values and protist taxa density in the blue network. Values for each taxon represent average body size taken from the literature. The figure shows that larger taxon Pca is more abundant in communities with higher averaged body size, while smaller taxa Tet and Chi are more abundance in communities with lower averaged body size. The different protist taxa are color coded and named by their abbreviations described in the Methods section.

**Figure 4.**
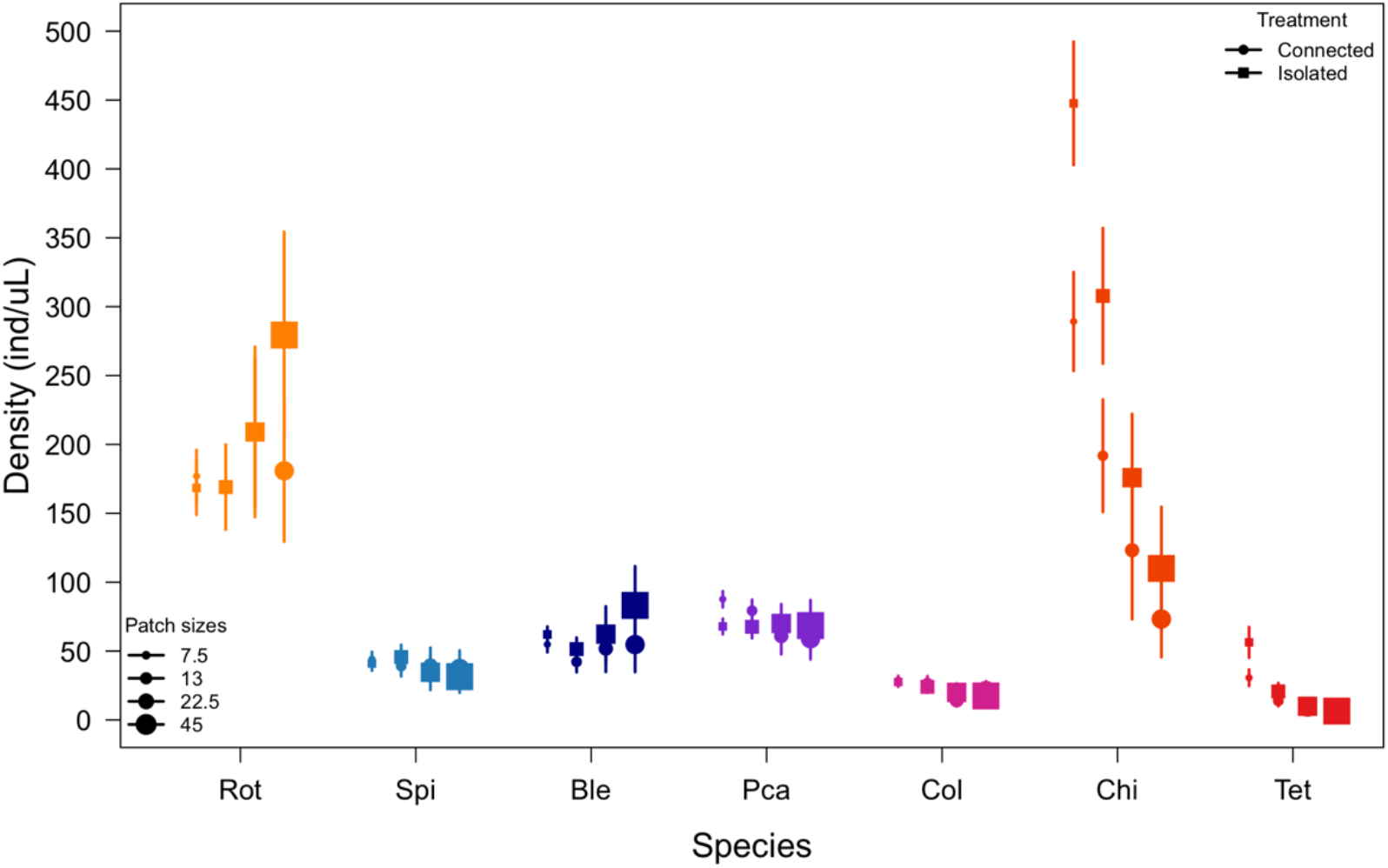
Effect of the cross-ecosystem resource pulse treatment on protist taxa density for the different ecosystem volumes (patch sizes) in the blue network. Each point represents the mean density and error bars represent the 95% confidence intervals. The figure shows that the effects of resource exchange on Ble, Pca, Chi and Tet shown in Figure 1 tend to disappear in larger patch sizes. The different protist taxa are color coded and named by their abbreviations described in the Methods section.

**Figure 5.**
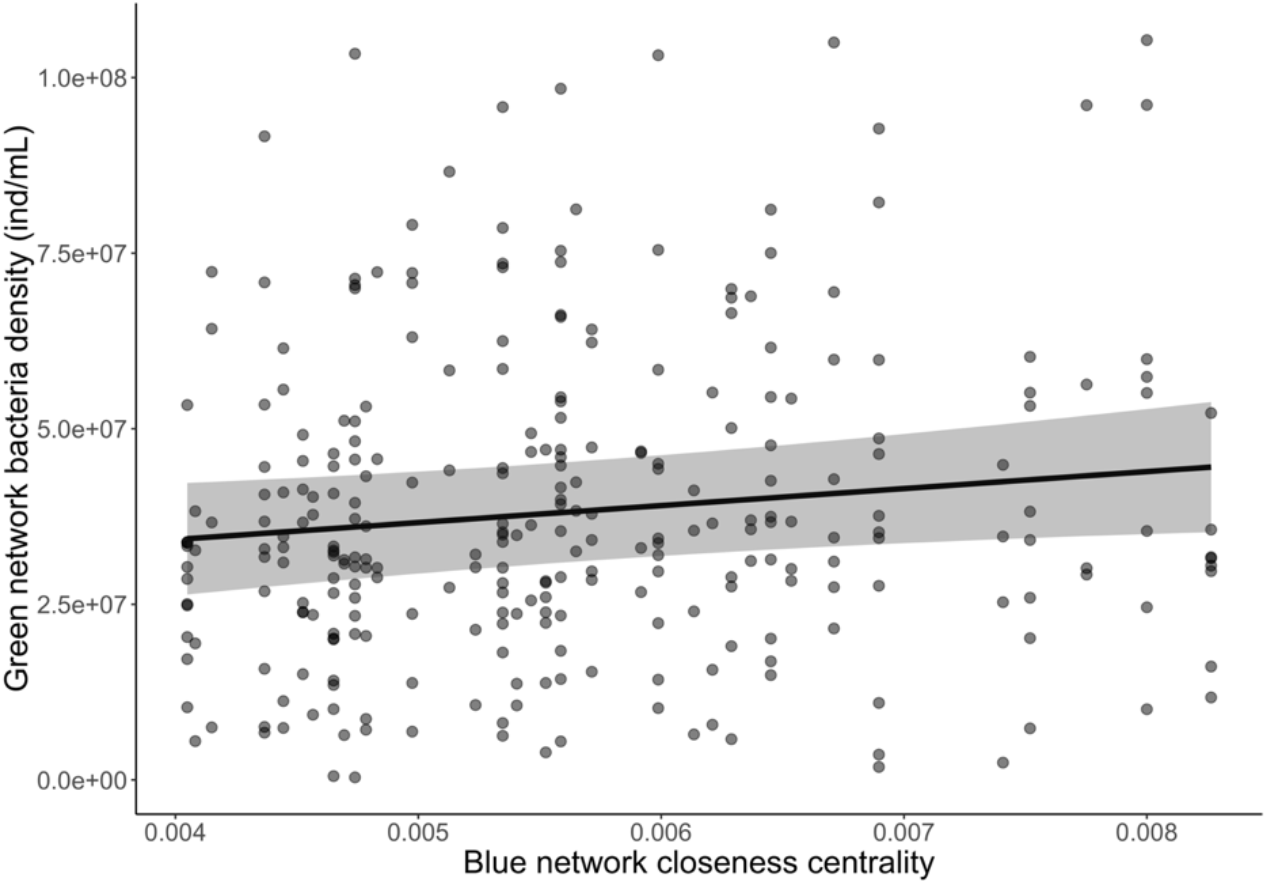
Effect of closeness centrality in the blue network on bacteria density in the connected green networks. Points represent the raw data. The black line is the prediction from the mixed effect model (Table S2), and shaded area represents the 95% confidence interval.

More specifically, in the blue network we found an increase in community mean but not in the standard deviation of trait values related to body size (mean body length and width, see Figure 2) when connected to the green network. Changes in community mean trait values resulted from increased population densities in the larger taxon Pca (1.2 times higher, CI=[1.09,1.29], see Figures 2 and 3) with resource exchange, accompanied by decreased population densities in the smaller taxa Tet (1.8 times lower, CI=[1.38, 2.25]), and Chi (1.6, CI=[1.35,1.77], see Figure 2 and 3).

Those variations in protist population densities with resource pulse led, at the community level, to an overall decline in total protist densities and richness and a marginal but significant increase in evenness in connected blue networks (Table S1), but with no detectable effect on total protist bioarea (a proxy for biomass, Table S1). We also found evidence that the effect of resource pulse in the blue network varied with ecosystem volume (‘R x V’ term in Table 1) with the differences between connected and isolated networks disappearing as ecosystem volume increases (Figure 4).

In the green network, total bacteria densities were 1.26 times (CI = [1.15, 1.37]) higher with than without resource pulse from the blue network (Table S1, Figure 2). This effect increased with closeness centrality in the blue network (Table S2, Figure 5).

## Discussion

The effect of cross-ecosystem resource exchange led to changes in protist population densities and varied depending on the specific spatial position of the cross-ecosystem coupling in the blue network. In the upstream ecosystems of the blue network, resource pulse from the green network favored one large taxon (*Paramecium*) but led to declines in the density of two smaller taxa (*Chilomonas*, *Tetrahymena*) compared to ecosystems in non-subsidized blue networks. In the downstream ecosystems, those effects were not detectable anymore, suggesting that the effect of cross-ecosystem resource exchange on populations diminished within the blue network in larger downstream positions. As hypothesized, we also observed a weak but significant spatial imprint (“mirroring” effect) of the blue network spatial structure on the green network. Total bacteria densities were higher in subsidized green ecosystems connected to more central ecosystems in the blue network. Consequently, localized cross-ecosystem fluxes, and their respective effects on recipient ecosystems, need to be studied from a perspective taking into account the explicit spatial configuration of the landscape.

Understanding why the signal of the mirroring effect from the blue to the green network in our experiment was observed but rather weak (see Table S2 and Figure 5) can lead to insights on the processes driving cross-ecosystem dynamics at landscape extent. Two fundamental elements from our experimental design could help to explain: i) dispersal and resource exchange happened at the same frequency (two times/week), and ii) the green lattice network has a total number of links much higher than in the blue dendritic network. In our experiment, the volume dispersed per edge was lower in the green (0.5 mL) relative to the blue (1 mL) network, however because the total number of links is higher in the green lattice network it would probably have taken a much lower dispersal volume to amplify the spatial signal and avoid homogenization. Our results suggest that dispersal needs to happen at a lower rate than resource exchange to cause a strong imprint of the connected network through spatial feedback, with the blue dendritic network imposing its own spatial constraints on the green lattice network. For metacommunities, it has been established that the balance between the speed of regional and local dynamics will drive their relative importance (e.g., mass effect vs. species sorting, Leibold *et al.* 2004; Leibold & Chase 2017). In metaecosystems, we propose that the balance between the speed of organism movement within a spatial network and of cross-ecosystem resource exchange might also be an important metric for expectations on the regional consequences of cross-ecosystem exchanges.

The effects of cross-ecosystem resource exchange on protist population densities in the blue network scaled up to affect aggregate community metrics leading to a decline in total protist densities and species richness in ecosystems connected to the green network. Generally, the effect of resource pulse is known to be destabilizing, affecting competitive outcomes leading to decreased richness and increased dominance by a few species (Stevens *et al.* 2004; Chase 2010; Cleland & Harpole 2010; Hautier *et al.* 2014). Interestingly, in our experiment, pulse of resource from the green ecosystem varied species relative abundance but did not change the dominance ranking in the protist communities in connected compared to isolated blue networks. For instance, in the smallest ecosystems, where the effects were the strongest, *Chilomonas* remained the dominant species despite being negatively affected by resource pulse (see Figure 4). Overall, the individual effects on each species population were strong enough to induce marginal effects on community evenness but not to shift species dominance between treatments.

The effects of cross-ecosystem resource exchange observed at the population level were still strong enough to alter community mean trait values related to body size in the community (see Figure 2). Those changes in mean trait values were associated to the density of specific taxa responding to resource exchange (see Figure 3). Cross-ecosystem fluxes selected for a larger taxa (Pca) and against the two smallest taxa (Tet, Chi). The balance between the one larger and less abundant taxon and the two smallest but more abundant taxa (see Figure 4), potentially explains why we did not observe any effect of resource pulse on protist bioarea (because of a cancelling-out differences between the two treatments). This change in community mean trait values also suggests that despite no observed change in bioarea (proxy for standing biomass) at the community level, cross-ecosystem resource exchanges can still likely affect ecosystem functions. For instance, in our experiment, subsidized ecosystems in the blue networks should have lower turnover rates due to larger individuals on average (Schramski *et al.* 2015) than non-subsidized ecosystems.

The relatively small effects of resource exchange observed in our experiment can be explained by different factors. Dispersal within the blue networks likely influenced the strength of the effect of localized cross-ecosystem resource exchange. It is well known that dispersal can prevent local extinctions (Hanski 1998). Especially, in our case, species that were negatively affected by cross-ecosystem resource pulse in upstream ecosystems of the blue network, might have been rescued by individuals dispersing from downstream ecosystems where the effect of resource pulse was not as strong. Eventually, this interaction between the effect of cross-ecosystem resource exchange and within-network dispersal constraints has significant and yet unexplored implications for the spatial re-arrangement of communities in the landscape.

Evolutionary history can also influence the effect of cross-ecosystem resource exchange on communities and ecosystems. A recent meta-analysis on the effect of cross-ecosystem subsidies showed that experimental studies in semi-artificial systems tend to have significantly lower effect sizes than observational studies (Montagano *et al.* 2019). One explanation for those results is that experimental studies are often conducted with organisms that have not necessarily evolved in an allochthonous resource pulse context (Holt 2008). Moreover, experimental studies in semi-artificial systems often connect systems that are very similar to one another, sharing the same evolutionary history (e.g., two protist communities), and thus also sharing similar traits (e.g., stoichiometry) making it less likely to observe spatial feedback (but see Gounand *et al.* 2017a).

Finally, our results have implications for natural systems because they suggest that upstream reaches might be more sensitive to terrestrial subsidy than larger downstream reaches and this spatial dependence in the strength of the effect can spatially feedback on connected ecosystems leading, at the landscape scale, to ‘mirrored’ dynamic and possibly functioning. Our experiment tested for the effect of landscape *per se*, all else being equals. In that context, differences in the effect of resource pulse were caused by intrinsic landscape characteristics: upstream patches had lower volumes and where less connected compared to larger volume and more connected downstream patches. Those attributes of dendritic networks were sufficient, despite the above discussed ecological and evolutionary weakening factors, to observe significant impacts on populations and mean community trait values, which are likely to also reflect changes in ecosystem functioning. In nature, however, all else is not equal. Upstream river patches are not only shallower and more isolated but they are also often more shaded, more heterotrophic and thus potentially more dependent on allochthonous subsidies than larger downstream patches (England & Rosemond 2004). Therefore, our results are conservative because in natural systems upstream patches are smaller and receive more subsidies, all else being equal.

Taken together, our results show how metaecosystem dynamics can impact the balance of community composition and trigger cross-ecosystem feedbacks that can spread across a whole landscape. Our study thus constitutes another illustration of the need to integrate spatial structure into land management, and confirm the need to incorporate more of the complexity of natural landscape into meta-ecosystem theory (Gounand *et al.* 2017b; Leroux *et al.* 2017).

## Acknowledgement

We thank M.-J. Fortin for kind discussions that led to the development of the analytical pipeline of this study. We also thank S. Gut, S. Flückiger and E. Keller for help during the laboratory work. Funding is from the Swiss National Science Foundation grants no. PP00P3_150698 and PP00P3_179089 (to FA), University of Zurich, Eawag, and the University of Zurich Forschungskredit (to IG). This is publication ISEM-YYYY-XXX of the Institut des Sciences de l’Evolution - Montpellier.

---

**Table S1.**
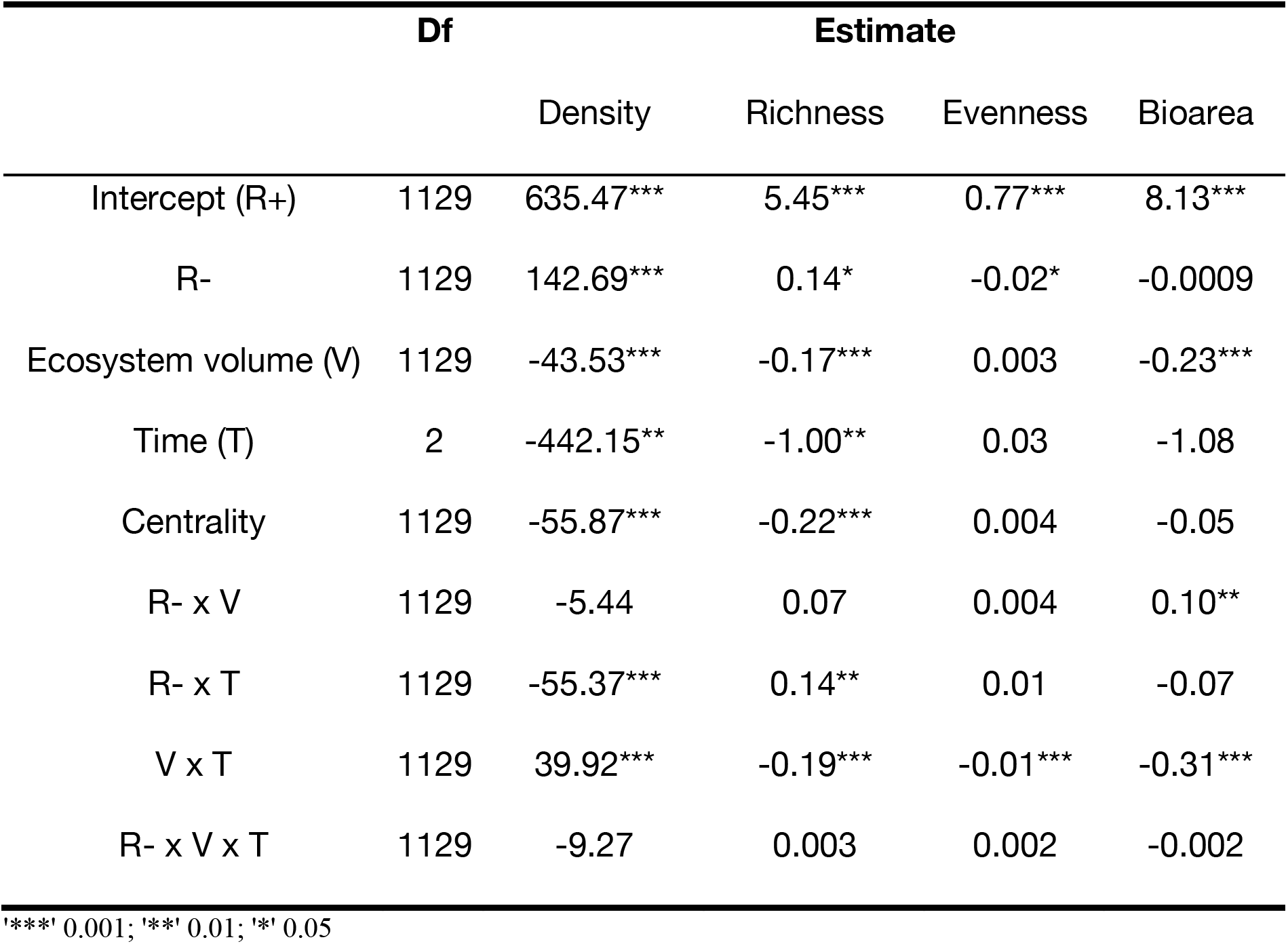
Mixed-effect models testing the effects of resource pulse on aggregate protist community metrics in the blue network: protist total density, richness, evenness and bio area per mL. The structure of the mixed-effect models is described in the Methods section. The fixed structure of each model (by column order) respectively explained 64%, 93%, 76%, and 29% of the total variance, the remaining going to replicates and experimental time variances. All numerical predictors were scaled. Protist bio area was log-transformed.

**Table S2.** Mixed-effect models testing the effects of blue network structure and properties on bacteria density (ind/mL) in the green network. The structure of the mixed-effect model is described in the Methods section. The fixed structure of the model explained 93% of the total variance, the remaining attributed to experimental time variance. All numerical predictors were scaled.

|  | **Df** | **Estimate** |
| --- | --- | --- |
| Intercept (R+) | 279 | 38.13e+06*** |
| Ecosystem volume | 279 | -1.61e+06 |
| Centrality | 279 | 3.95e+06* |
| Protist density | 279 | -0.36e+06 |
| Protist richness | 279 | 0.35e+06 |
| Protist bioarea | 279 | 2.45e+06 |

\*\*\*' 0.001; '\*\*' 0.01; '\*' 0.05

